# Fractional Dosing of Yellow Fever Vaccine to Extend Supply: A Modeling Study

**DOI:** 10.1101/053421

**Authors:** Joseph T. Wu, Corey M. Peak, Gabriel M. Leung, Marc Lipsitch

**Affiliations:** WHO Collaborating Centre for Infectious Disease Epidemiology and Control, School of Public Health, Li Ka Shing Faculty of Medicine, The University of Hong Kong, Hong Kong Special Administrative Region, China; Center for Communicable Disease Dynamics, Department of Epidemiology, Harvard T.H. Chan School of Public Health, 677 Huntington Avenue, Boston MA 02115 USA; Department of Immunology and Infectious Diseases, Harvard T.H. Chan School of Public Health, 665 Huntington Avenue, Boston MA 02115 USA.

## Abstract

**Background:** The ongoing yellow fever (YF) epidemic in Angola strains the global vaccine supply, prompting WHO to adopt dose sparing for its vaccination campaign in Kinshasa in July-August 2016. Although a 5-fold fractional-dose vaccine is similar to standard-dose vaccine in safety and immunogenicity, efficacy is untested. There is an urgent need to ensure the robustness of fractional-dose vaccination by elucidating the conditions under which dose fractionation would reduce transmission.

**Methods:** We estimate the effective reproductive number for YF in Angola using disease natural history and case report data. With simple mathematical models of YF transmission, we calculate the infection attack rate (IAR, the proportion of population infected over the course of an epidemic) under varying levels of transmissibility and five-fold fractional-dose vaccine efficacy for two vaccination scenarios: (i) random vaccination in a hypothetical population that is completely susceptible; (ii) the Kinshasa vaccination campaign in July-August 2016 with different age cutoff for fractional-dose vaccines.

**Findings:** We estimate the effective reproductive number early in the Angola outbreak was between 5·2 and 7·1. If vaccine action is all-or-nothing (i.e. a proportion VE of vaccinees receives complete and the remainder receive no protection), *n*-fold fractionation can dramatically reduce IAR as long as efficacy VE exceeds 1/*n*. This benefit threshold becomes more stringent if vaccine action is leaky (i.e. the susceptibility of each vaccinee is reduced by a factor that is equal to the vaccine efficacy VE). The age cutoff for fractional-dose vaccines chosen by the WHO for the Kinshasa vaccination campaign (namely, 2 years) provides the largest reduction in IAR if the efficacy of five-fold fractional-dose vaccines exceeds 20%.

**Interpretation:** Dose fractionation is a very effective strategy for reducing infection attack rate that would be robust with a large margin for error in case fractional-dose VE is lower than expected.

**Funding:** NIH-MIDAS, HMRF-Hong Kong

## INTRODUCTION

Yellow fever (YF) has resurged in Angola and threatens to spread to other countries with relatively low YF vaccine coverage. As of 8 July 2016, YF cases have been exported from Angola to Kenya (2 cases), China (11), and DRC (59), raising concern that YF could resurge in other populations where competent vectors are present and vaccine coverage is low.^1,2^ Indeed, DRC has already declared a YF epidemic in Kinshasa and two other provinces. A broad band of sub-Saharan Africa north of Namibia and Zambia is at risk (http://www.cdc.gov/yellowfever/maps/africa.html), as is much of the northern portion of South America (http://www.cdc.gov/yellowfever/maps/south_america.html). The global community is increasingly concerned for the risk of YF emergence in Asia, where the disease has been curiously absent despite seemingly amenable conditions.

There is a safe, highly effective live-attenuated vaccine against YF.^3^ However, the global emergency stockpile of YF vaccines, which has been maintained at approximately 6·8 million doses before 2016, has already been depleted twice by the Angola outbreak. With a throughput of only 2 to 4 million doses per month, YF vaccine supply is inadequate given the large urban populations at risk for YF infection. In response to such shortage, dose fractionation has been proposed to maximize the public health benefit of the available YF vaccines.^4^ Under dose fractionation, a smaller amount of antigen would be used per dose in order to increase the number of persons who can be vaccinated with a given quantity of vaccine.^3^ This strategy was previously proposed to extend pre-pandemic influenza vaccine supplies.^5^ If dose fractionation were consistently adopted, equity of YF vaccine access would also be enhanced both within and across countries at risk, as more people could benefit from vaccination without depriving others.^6^

Indeed, following the SAGE endorsement on 17 June 2016, the WHO recommended dose fractionation in its emergency YF vaccination campaign in July-August 2016 to vaccinate 8 million people in Kinshasa, 3 million in anterior Angola and 4·3 million along the DRC-Angola corridor.^7^ Specifically, 2·5 million standard-dose vaccines would be allocated to Kinshasa where 200,000 standard-dose vaccines would be given to children age 9 months to 2 years and the remaining allocation are to be fractionated five-fold and administered to the rest of the population.

The evidence base for fractional-dose YF vaccines is built upon two studies that compared the safety and immunogenicity of standard-dose and five-fold fractional-dose YF vaccines. The first is a randomized, noninferiority trial which showed that 0·1 ml intradermal (ID) vaccination with the 17D YF vaccine was equally safe and immunogenic compared to the standard 0.5ml subcutaneous vaccination.^8^ The second is a randomized trial of subcutaneous administration of the 17DD vaccine given in Brazil which showed that there was no significant difference in immunogenicity and viremia kinetics when the currently administered vaccine (containing 27,476 IU of virus) was given at subdoses as low as 11% of the full dose (3,013 IU).^9^ Even lower doses produced noninferior immune responses, but not equivalent viremia kinetics.^9^ For comparison, the WHO minimum for YF vaccines is 1,000 IU per dose at the end of shelf life.^10^

No efficacy trial of YF vaccines, however, has ever been performed in humans,^11^ so the comparative efficacy of different doses and routes of administration remains uncertain. In particular, it is not known whether equal immunogenicity implies equal vaccine efficacy for YF vaccines. Moreover, the findings of equal immunogenicity of reduced doses are limited to healthy adults; no comparable data exist in children (thus the age cutoff of 2 years for fractional-dose vaccines in Kinshasa), elderly or immunocompromised individuals (e.g. HIV-infected people, pregnant women, etc.). As such, while noninferior immunogenicity of fractional-dose vaccines provide a strong basis for an initial consideration of dose-sparing strategies for YF vaccines, it would be prudent to ensure the robustness of this strategy by carefully evaluating the risk and epidemiologic impact of reduced vaccine efficacy in fractional-dose vaccines. Such an evaluation is nontrivial because even if dose fractionation reduces vaccine efficacy, higher vaccine coverage may confer higher herd immunity in which case the number of infections could be significantly reduced by the indirect effect of large-scale vaccination.^12^ The lower the transmissibility, the larger the number of infections that can be averted by indirect protection, as illustrated by the previous study of dose fractionation for prepandemic influenza vaccines.^5^ The importance of herd immunity for YF vaccination is unknown because transmissibility of YF in urban settings has never been adequately characterized due to limited data.

To strengthen the evidence base for the public health benefit of dose fractionation of YF vaccines, we use simple mathematical models to assess the potential reduction in infection attack rate (IAR, defined as the proportion of population infected over the course of a sustained epidemic) conferred by five-fold dose fractionation under different epidemic scenarios and reductions in vaccine efficacy. We find that all dose-sparing strategies considered are likely to provide significant benefit epidemiologically, and that the best policy will be determined by balancing logistical and regulatory considerations against the extent of epidemiologic benefit. In particular, we conclude that the WHO Kinshasa dose-sparing vaccination campaign in July-August 2006 would be an effective strategy for reducing infection attack rate, and the results would be robust against a large margin for error in case five-fold fractional-dose efficacy turns out to be lower than expected.

## METHODS

### Estimating the epidemiologic parameters for YF

First, to parameterize realistic epidemic scenarios for our analysis, we estimate the reproductive number of YF over the course of the Angola outbreak and use the estimates during the early epidemic stages (before large-scale vaccination affected transmission) as the range of basic reproductive number (R_0_) for future outbreaks in other populations. To this end, we use the Wallinga and Teunis method^13^ to estimate the reproductive number of YF from the daily number of confirmed YF cases recorded in the 17 April 2016 WHO Angola Situation Report,^14^ assuming that all cases were attributed to local transmission (i.e. no importation of cases). When estimating the extrinsic incubation period, we assume that the average temperature in Angola was 28 degrees Celsius during the outbreak. To estimate the serial interval distribution, we make the following assumptions: (i) the extrinsic incubation period follows the Weibull distribution estimated by ref. ^15^ which has mean 12·7 days at 28 degrees Celsius; (ii) the intrinsic incubation period follows the lognormal distribution estimated by ref. ^15^ which has mean 4·6 days; (iii) the infectious period in human is exponentially distributed with mean 4 days;^16^ (iv) the mosquito lifespan is exponentially distributed with mean 7 to 14 days.^17^ We estimate the initial reproductive number of the YF outbreak in Angola as the average reproductive number among all cases who developed symptoms one serial interval before vaccination campaign began to affect disease transmission (see Figure 1).

**Figure 1:**
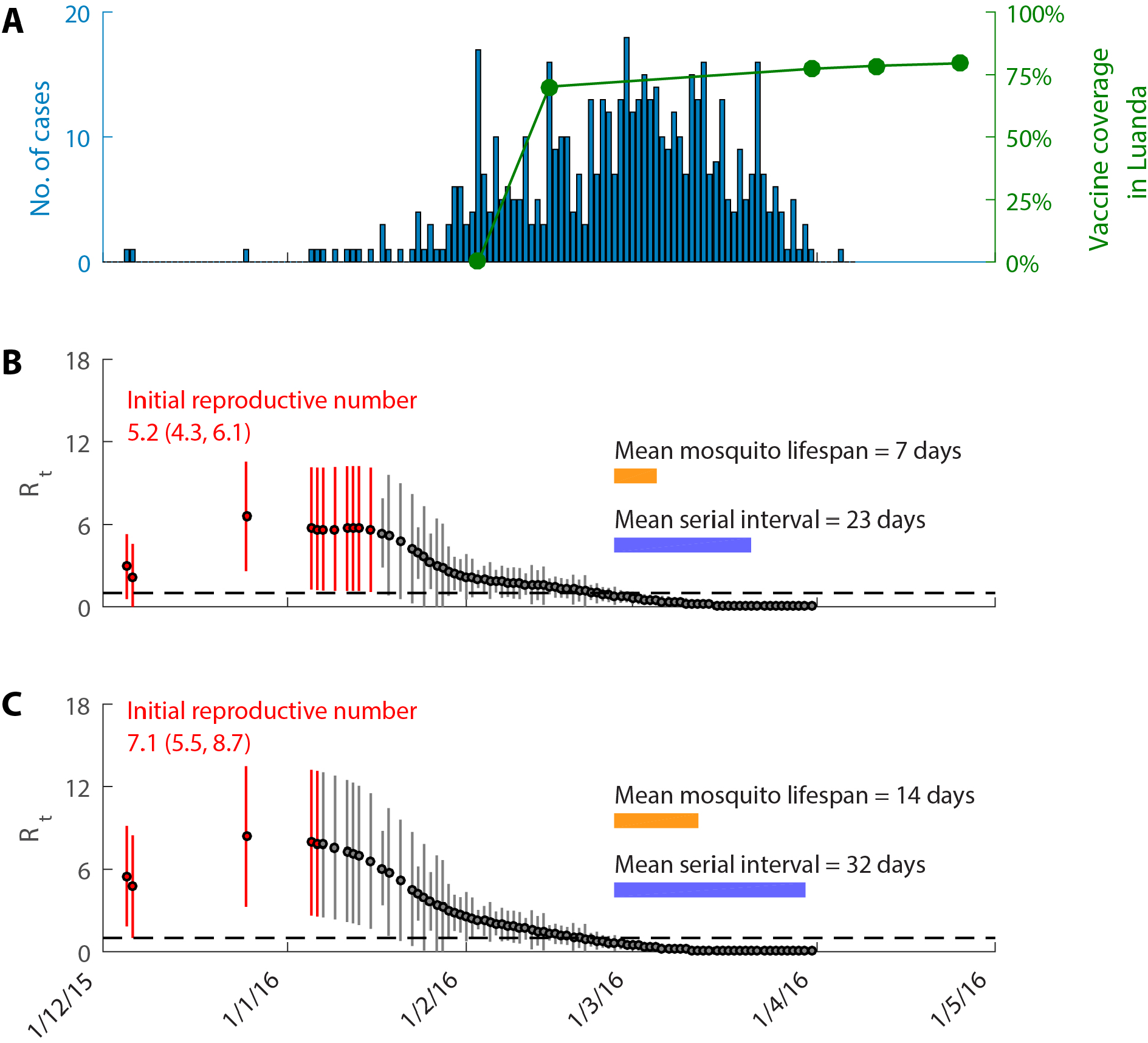
Estimates of reproductive number over the course of the Angola epidemic. **A.** Epidemic curve of confirmed cases by dates of symptom onset in Angola and vaccine coverage in Luanda province achieved by the reactive YF vaccination campaign that started on 2 February 2016.^28^ The first cases of this YF outbreak were identified in Luanda province which accounted for 90 of the 121 cases confirmed in Angola up to 26 February 2016. **B-C** Estimates of the daily reproductive number (*R*_t_) assuming that the mean mosquito lifespan was 7 and 14 days, respectively. The red data points correspond to the cases that were used to estimate the initial reproductive number. These cases had symptom onset one mean serial interval before the vaccination campaign began to affect disease transmission (which was assumed to be 7 days after the start of the campaign to account for the time it takes for adaptive immunity to develop). The orange and purple horizontal bars indicate the length of the mean mosquito lifespan and serial interval on the scale of the x-axis, respectively.

### Dose-response for fractional-dose vaccines

Let *S*_0_ be the proportion of population susceptible just before the vaccination campaign begins and *V* be the vaccine coverage achievable with standard-dose vaccines. Suppose each standard-dose vaccine can be fractionated into *n, n*-fold fractional-dose vaccines (i.e. each of which contains 1/*n*-th the amount of the antigen in a standard-dose vaccine) with vaccine efficacy *VE*(*n*). That is, the vaccine efficacy of standard-dose vaccines is *VE*(1) which was assumed to be 1. Given *V*, the highest fractionation sensible is *n*_max_ = *S*_0_/*V* if the susceptible population can be identified for targeted vaccination and *n*_max_ = 1/*V* otherwise, i.e. the fractionation *n* must lie between 1 and *n*_max_. To avoid overstating the benefit of dose fractionation, we assume that vaccine efficacy of *n*-fold fractional-dose vaccines for *n* between 1 and 5 increases linearly with the amount of antigen in the vaccines (see appendix for explanation). Potential increases in vaccine wastage during dose-sparing would be mostly due to unused, reconstituted vaccines^18^ or increased vaccine failure due to inexperience with intradermal administration among vaccinators. In the setting of mass vaccination campaigns, wastage due to unused vaccine doses will likely to be negligible because vaccination sessions will be large.

### Infection attack rate

We use IAR as the outcome measure for evaluating the impact of dose fractionation. We calculate IAR using the classical final size approach which is exact for directly transmitted SIR-type diseases^19^ but only an approximation for vector-borne diseases.^20^ Nonetheless, this approximation is excellent over realistic parameter ranges because only a very small proportion of mosquitoes are infected with YF virus even during epidemics (necessitating pooled testing).^21^ See appendix for the mathematical details.

We denote the IAR under *n*-fold dose fractionation by *IAR*(*n*). To evaluate the outcome of fractional-dose vaccination against that of standard-dose vaccination, we calculate the absolute and relative reductions in IAR as *IAR*(1) – *IAR(n)* and 1 – *IAR(n)/ IAR(1)*, respectively. We assume that the vaccination campaign is completed before the start of the epidemic.

### Vaccine action

We assume that vaccine action is all-or-nothing, i.e. *n*-fold fractional-dose vaccines provide 100% protection against infection in a proportion *VE*(*n*) of vaccinees and no protection in the remainder. In this case, *n*-fold dose fractionation results in lower IAR if and only if the vaccine efficacy of *n*-fold fractional-dose vaccines is at least 1/*n* times that of standard-dose vaccines, i.e. *VE*(*n*) > *VE*(1)/*n* (see appendix for details). We term this the benefit threshold for dose fractionation. We also consider the alternative case in which vaccine action is leaky, i.e. *n*-fold fractional-dose vaccines reduce the hazard of infection (the probability of disease transmission per mosquito bite) of each vaccinee by a proportion *VE*(*n*).^22,23^ Compared to all-or-nothing vaccines, leaky vaccines have substantially higher benefit thresholds, especially when transmissibility is high (see Results). However, YF vaccine action is much more likely to be all-or-nothing than leaky (see Discussion). As such, we present our main results in the context of all-or-nothing vaccine action. In principle, disease transmission can be halted if the effective vaccine coverage, defined as the proportion of population immunized (e.g.*V×n×VE*(*n*) if vaccination comprises *n*-fold fractional-dose vaccines only), exceeds the herd immunity threshold 1 −1/*R*_0_.

### Vaccination scenarios

We consider two vaccination scenarios with various levels of transmissibility and efficacy reduction in fractional-dose vaccines:

1. *Random vaccination in a hypothetical population*. To illustrate the potential impact of dose fractionation, we first consider a hypothetical scenario where the entire population is susceptible (*S*_0_ = 1) and each individual has the same probability of receiving vaccination. We compare the outcome of using the entire vaccine stockpile for either standard-dose or five-fold fractional-dose vaccination. If some individuals are immune (*S*_0_ < 1, due to previous exposure or vaccination) and vaccination can be targeted at susceptible individuals only, then the resulting IAR within the susceptible population would be the same as that for random vaccination in a completely susceptible population with coverage *V*/ *S*_0_ and basic reproductive number *R*_0_*S*_0_.
2. *The Kinshasa vaccination campaign in July-August 2016*. The population size of Kinshasa is estimated to be around 12.46 million (https://www.cia.gov/library/publications/the-world-factbook/graphics/population/CG_popgraph%202015.bmp) and around 2·5 million standard-dose vaccines is expected to be administered to the Kinshasa population during this vaccination campaign (personal communication, Bruce Aylward and Alejandro Costa, WHO). Under the WHO dose-sparing strategy, 200,000 standard-dose vaccines will be given to children aged between 9 months and 2 years (which is sufficient for vaccinating all unvaccinated children in this age range) and the remaining allocation will be fractionated to one-fifth of the standard dose and given to the rest of the population. We compare the outcome when the vaccines are administered (i) in standard dose only (strategy S) and (ii) according to the WHO dose-sparing strategy with alternative age cutoffs for fractional-dose vaccines ranging from 2 to 20 years (strategy F). For the latter, let *Z* be the age cutoff and *p*(*Z*) be the proportion of population targeted for standard-dose vaccination. For a given standard-dose vaccine coverage *V*, the proportion of population receiving standard-dose and fractional-dose vaccines are min(*V*, *p*(*Z*)) and 5max(*V* − *p*(*Z*),0), respectively. Therefore, the effective vaccine coverage after the vaccination campaign is *B* + min(*V, p*(*Z*))· *VE*(1) + 5max(*V* − *p*(*Z*),0)· *VE*(*5*) where *B* is the vaccine coverage immediately before the campaign (i.e. at the end of June 2016). See appendix for the calculation details.

## RESULTS

### Reproductive number of yellow fever in Angola

Figure 1 shows that the initial reproductive number of YF in Angola was 5·2 (95% CI 4·3, 6·1) and 7·1 (5·5, 8·7) if the mean mosquito lifespan was 7 and 14 days, respectively. While these estimates may reflect partial immunity due to prior vaccination or exposure among some of the population (we estimated that around 28% of the Angola population had been vaccinated before the YF epidemic; see appendix for calculation details), we assume that the basic reproductive number of a future outbreak in another population would range between 4 and 12 due to varying vector ecology and levels of preexisting immunity in the population.

### Random vaccination in a hypothetical population

Figure 2A–B shows the effective vaccine coverage and IAR under standard-dose and fractional-dose vaccination as a function of standard-dose vaccine coverage *V* given varying levels of transmission and five-fold fractionation vaccine efficacy when vaccine action is all-or-nothing. Figures 2C–D show the corresponding absolute and relative reduction in IAR and confirm our earlier claim that fractional-dose vaccination reduces IAR when *VE*(5) > *VE*(1)/ 5 = 0.2. Fractional-dose vaccination substantially reduces IAR if *V* > 10% and such reduction only diminishes to insignificant levels when *V* is close to the herd immunity threshold 1 −1/*R*_0_ (e.g. 75% and 88% for *R*_0_ = 4 and 8, respectively). In short, dose fractionation reduces IAR when (i) the standard-dose vaccine supply is insufficient to halt disease transmission and (ii) fractional-dose vaccine efficacy is above 0·2.

**Figure 2:**
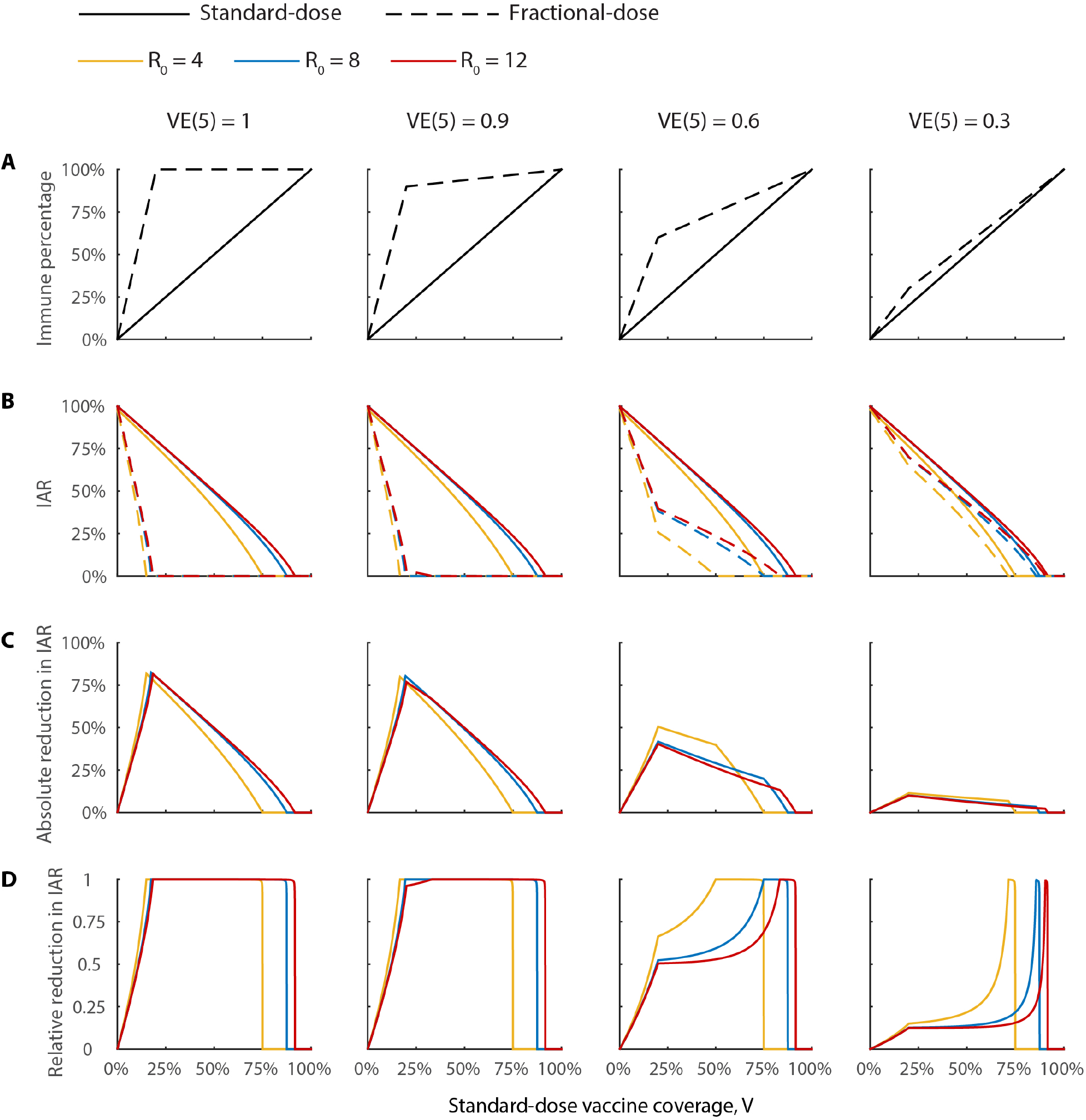
The impact of five-fold fractional-dose vaccination with different vaccine efficacy and reproductive numbers. We assume that (i) the whole population is susceptible, (ii) vaccine action is all-or-nothing, and (iii) standard-dose vaccine efficacy is 1. If the standard-dose vaccine coverage *V* exceeds 20%, then everyone in the population can be vaccinated under five-fold fractionated-dose vaccination, in which case the fractionation would only be *n* = 1/*V*. A The effective vaccine coverage (*VE*(*n*) × *nV*), which is essentially the percentage of population immunized, as a function of standard-dose vaccine coverage *V* under standard-dose vaccination (solid curves) and five-fold fractional-dose vaccination (dashed curves). **B** Infection attack rate (IAR) under standard-dose vaccination and five-fold fractional-dose vaccination. IAR is reduced to 0 when the effective vaccine coverage reaches the herd immunity threshold 1 −1/*R*_0_. **C** Absolute reduction in IAR. As *V* increases from 0, a kink appears when the herd-immunity threshold is attained or everyone is vaccinated under five-fold fractional-dose vaccination (i.e., *V* = 20%). If five-fold fractional-dose vaccination at 100% coverage cannot attain the herd immunity threshold (because of low fractional-dose vaccine efficacy), then a second kink appears when V is large enough such that fractional-dose vaccination attains herd-immunity threshold due to the increase in *VE*(*n*) resulting from lower fractionation (namely *n* = 1/*V*). **D** Relative reduction in IAR.

If vaccine action is “leaky,” then the benefit threshold (the efficacy of *n*-fold fractionated doses necessary to reduce IAR) is higher than 1/*n* and increases with transmission intensity (Figure 3). This occurs because under the leaky model each infectious bite is assumed to be less likely to cause infection if the host is vaccinated, but the probability of infection grows as the person receives more infectious bites. Figure 3 shows, under the leaky model of vaccine action, dose fractionation is much less beneficial if vaccine action is leaky, efficacy is modest, and R_0_ is high. See appendix for the mathematical details.

**Figure 3:**
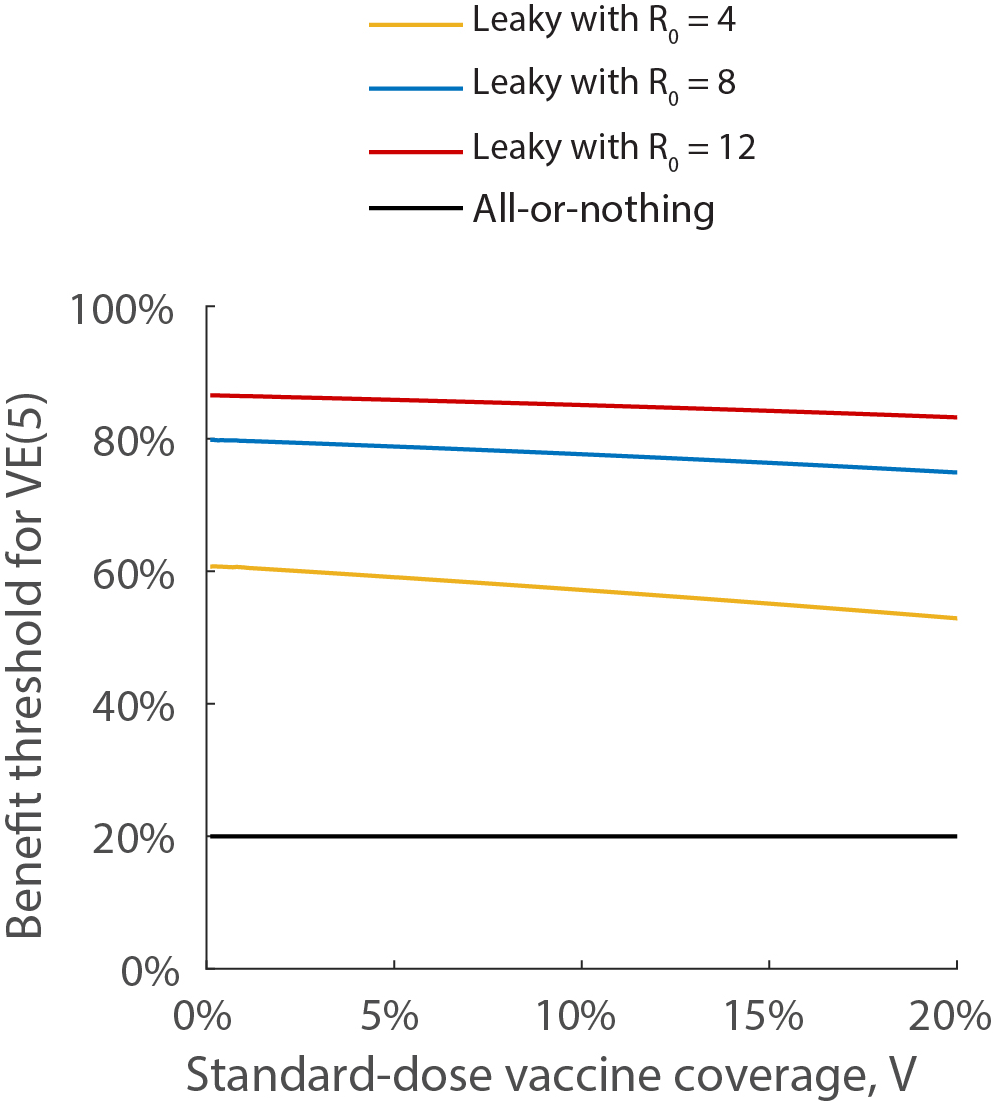
Benefit thresholds for leaky vaccines as a function of standard dose vaccine supply *V* and basic reproductive number *R*_0_. Five-fold fractionated dosing reduces IAR compared to standard dosing if the leaky vaccine efficacy of fractional-dose is above the threshold. This threshold becomes high for large values of *R*_0_ because under the “leaky” vaccine action model, multiple exposures eventually overcome vaccine protection and lead to infection of vaccinated individuals.

**Figure 4.**
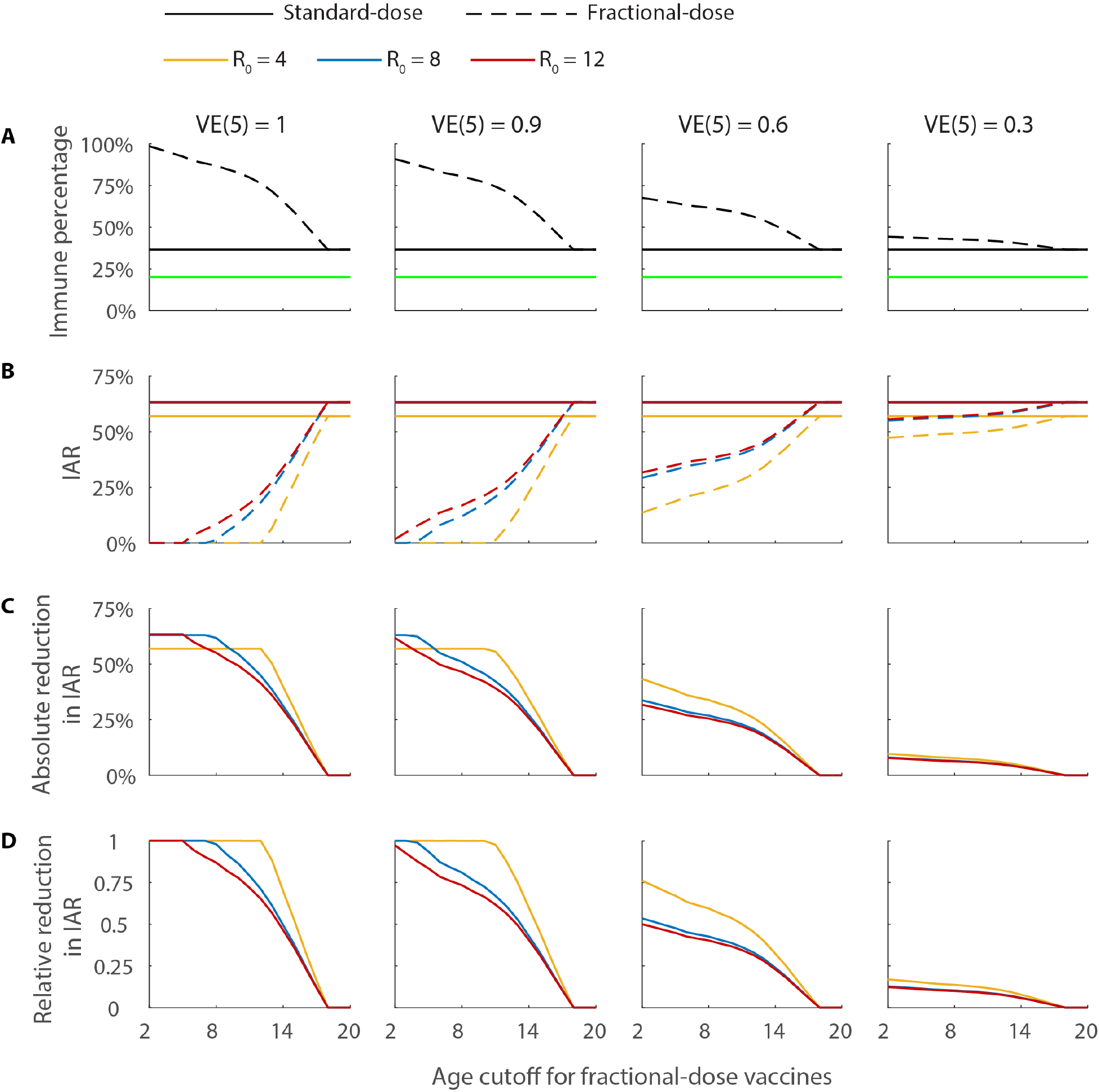
Evaluating the WHO dose-sparing strategy in the Kinshasa vaccination campaign in July-August 2016. **A** Effective vaccine coverage (i.e. the percentage of population immunized). The green line indicates the pre-campaign coverage. The black solid and dashed lines indicate the post-campaign coverage if vaccines are administered (i) in standard dose only (strategy S) and (ii) according to the WHO dose-sparing strategy with alternative age cutoffs for fractional-dose vaccines (strategy F). **B** IAR under strategy S and F for different R_0_. **C-D** Absolute and relative reduction in IAR.

A recent study suggested that the mosquito biting rate for individuals aged 20 or above is 1·22 times higher than those age under 20.^24^ We performed a sensitivity analysis to show that our results are unaffected by such heterogeneity. See “Hetereogeneity in biting rates” in the appendix for details.

### The WHO vaccination campaign in Kinshasa

We estimate that the vaccine coverage in Kinshasa was 20% at the end of June 2016 before the vaccination campaign began. The WHO vaccination campaign would increase the effective vaccine coverage to 37% if all the vaccines were administered only in standard dose. Under the WHO dose-sparing strategy, the effective vaccine coverage can be increased to 99%, 91%, 68% and 44% if the vaccine efficacy of five-fold fractional-dose vaccines VE(5) is 1, 0·9, 0·6 and 0·3, respectively. The corresponding absolute reduction in IAR is 57%, 57%, 43% and 10% if *R*_0_ = 4 and around 63%, 63%, 32% and 8% if *R*_0_ is 8 to 12. These IAR reductions correspond to 7·10, 7·10, 5·36, 1·25 million infections averted if *R*_0_ = 4 and around 7·85, 7·85, 3·99 and 1·0 million infections averted if *R*_0_ is 8 to 12. The age cutoff for fractional-dose vaccines chosen by the WHO (namely, 2 years) provides the largest reduction in IAR as long as VE(5) is above 0·2. That is, the WHO dose-sparing strategy is optimal as long as five-fold fractional vaccines are at least 20% efficacious. These figures are based on the assumption of a sustained epidemic such that transmission declines when the population of susceptible hosts is depleted.

## DISCUSSION

Our primary analysis shows that dose fractionation of YF vaccine, if there is no loss of efficacy as currently assumed, could provide a substantial benefit to reducing the attack rate of YF in a population. We consider this assumption of full efficacy for five-fold fractionation to be the most likely scenario, despite the lack of efficacy data on any YF vaccine, for several reasons: 1) two studies of five- or greater-fold vaccination doses have shown indistinguishable immunogenicity in humans; 2) at least some preparations of YF vaccine substantially exceed the WHO minimum standard for potency of 1,000 IU/dose, so fractionation at some level could be performed without dropping below that threshold; 3) YF vaccine is live attenuated virus, so a biological rationale exists that if a productive vaccine-virus infection can be established by a fractionated dose, protection should be comparable to that with a higher dose. Nonetheless, to assess the robustness of the conclusion that dose fractionation is likely to be beneficial, against the possibility that in fact efficacy of fractionated doses is lower than anticipated, we consider the possibility that five-fold fractionated dosing fails to immunize a proportion (1-*VE*(5)) of recipients. We find that as long as at least 20% of recipients are fully immunized by the vaccine, more people would be immunized by vaccinating five times as many people with one-fifth the dose, and so the population-wide benefits of higher coverage would outweigh the lower efficacy of fractionated dosing for individual vaccinees.

Even more unlikely, in our opinion, is that fractionated doses would be substantially less efficacious according to a “leaky” model, in which all vaccinated individuals were imperfectly protected against infection from each infectious bite, with the same probability of infection from each bite, reduced by vaccine by a proportion VE (see appendix for details). If this were the case however, we found that especially in high-transmission areas, the fractionated-dose vaccine would need to be 80–90% efficacious to provide a benefit over standard dosing.

Our analysis is not intended to recommend extending coverage to the point of knowingly compromising efficacy. Rather, our analysis indicates that a strategy of fractionation to a dose that provides equivalent immunogenicity to standard dosing would be greatly beneficial if efficacy is equivalent to standard dosing, and would still be beneficial if, unexpectedly, efficacy were somewhat lower than for standard dosing.

We have used five-fold fractionation as an example because it is the strategy with the best evidence base of equal immunogenicity. However, some data suggest that more than five-fold fractionation could be equally immunogenic, and of course the benefits of fractionation would be greater if more than five-fold fractionation were logistically possible and comparably efficacious.

We have considered fractional dosing for residents of areas at high risk for transmission. Another group of interest are travelers, for whom we must also consider longevity of response, lower levels of exposure, and more detailed discussions on equity outside the scope of this modeling paper. The cost of fractional-dose strategies will depend on the route of administration, but could potentially be substantially less expensive per vaccine recipient.^18^

Our simple model has several limitations. We assume homogeneous mixing of the population (reasonable at least locally for a vector-borne disease). We also fix a particular value of *R*_0_ for each calculation, and assume this value is maintained until the epidemic has swept through a population. In reality, R_0_ will vary seasonally as vector abundance, extrinsic incubation period, and other factors vary. The existence of a high-transmission season might enhance the benefits of fractional-dose vaccination. Most importantly, there will be a premium on achieving high vaccine coverage before the peak of transmission to maximally impact transmission, and this will be limited by supply constraints that could be partially relieved by fractionation. However, the cases-averted estimates might not all be achieved in a single transmission season if seasonal declines in mosquito abundance abrogate transmission before the large majority of the population has become infected.

We have focused on the benefits of increasing vaccine coverage within a single population. Given the global shortage of YF vaccines, an additional benefit of fractionated dosing is to extend coverage to a wider geographic area, covering more populations with vaccine than could be achieved with standard dosing. Indeed, part of the WHO plan is to vaccinate border areas between Angola and Congo^25^, providing benefit to that population as well as an “immune buffer” to slow movement of disease toward Kinshasa.^26^

We conclude that dose fractionation could be a very effective strategy for improving coverage of YF vaccines and reducing infection attack rate in populations -- possibly by a large absolute and relative margin -- if high to moderate efficacy is maintained by reduced-dose formulations. For vaccines whose standard formulations exceed WHO minimum concentration of viral particles,^10^ this dose fractionation could be accomplished without changing the WHO recommendations. In particular, the WHO plan to use fractional dosing to extend the coverage of vaccination within Kinshasa and in surrounding areas is robust in the sense that it is expected to provide greater benefit than the use of full dosages, even if, counter to current evidence, efficacy of fractionated doses is substantially lower than that of standard doses.

Rollout of fractionated dosing should perhaps be preceded or accompanied by noninferiority studies of the intended vaccine’s immunogenicity at fractional doses in the intended populations. Ongoing programs should be monitored by observational studies of safety, immunogenicity and, if possible, effectiveness^18^ to assure that the assumptions underlying the rationale for such programs continue to be met. However, it is worth noting that if full-dose vaccine efficacy is indeed 100% or nearly so as currently believed, estimating the relative efficacy of fractional vs. standard doses in a comparative study would be challenging or impossible, as there might be few or no cases accrued in the standard-dose arm.

## Contributors

JTW, CMP, and ML reviewed the literature and designed the study. JTW and ML developed the mathematical model. JTW ran the mathematical model. JTW, CMP, GML, and ML interpreted the model results and approved the final version.

## Acknowledgements

We thank Jack Woodall and Rebecca Grais for comments on an earlier draft and Alejandro Costa and the students of EPI 260 for helpful discussions. This study was funded by Cooperative Agreement U54GM088558 from the National Institute Of General Medical Sciences, National Research Service Award T32AI007535–16A1, and a commissioned grant from the Health and Medical Research Fund from the Government of the Hong Kong Special Administrative Region. The content is solely the responsibility of the authors and does not necessarily represent the official views of the National Institute Of General Medical Sciences or the National Institutes of Health.

## Panel: Research in context

### Systematic review

We searched PubMed and Google Scholar on June 10, 2016, with the terms “yellow fever” and “vaccine” or “dose sparing”. We did not find any reports of randomized trials of yellow fever (YF) vaccine efficacy, at full or lower doses. Three relatively recent studies suggest similar immunological responses at five-fold, or more, fractionation as compared to the current dose antigen levels.^8,9,27^ While several recent perspective articles propose the dose-sparing strategy in response to the current shortage,^2–4^ to our knowledge this is the first study to test the robustness of this strategy (in terms of its epidemiologic impact) against the uncertainties surrounding fractional-dose YF vaccine efficacy and mode of action (e.g. “all-or-nothing” and “leaky”).

### Added value of the study

We estimate that the reproductive number of the YF epidemic in Angola during 2016 is 5·2 to 7·1. Our study is the first to provide an estimate of the transmissibility of YF in urban settings. We characterize the threshold of vaccine efficacy above which dose-sparing can drastically reduce the number of YF cases. We show how the benefits of dose fractionation are influenced by the transmission intensity of the setting, the target coverage, and the fractional-dose vaccine efficacy and mode of action. We show that the WHO dose-sparing strategy for the Kinshasa vaccination campaign in July-August 2006 is a robust and effective strategy for reducing infection attack rate that would be robust with a large margin for error in case five-fold fractional-dose vaccine efficacy turns out to be lower than expected.

### Interpretation

Our results support the growing evidence that dose-sparing strategies should be adopted as an option for extending the currently sparse YF vaccine supply.

## Appendix

## Estimation of the effective reproductive number for YF in Angola

We use the Wallinga and Teunis method^13^ to estimate the reproductive number over the course of the YF outbreak in Angola from the daily number of confirmed cases recorded in the 17 April 2016 WHO Angola Situation Report.^14^ We assume that all cases were attributed by local transmission, i.e. no importation of cases. Let *t_i_* be the date of symptom onset for case *i*. The relative likelihood that case *i* has been infected by case *j* is

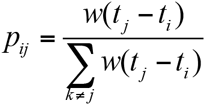

where *w*(·) is the probability density function of the serial interval. Assuming that the probability of case *j* infecting case *i* is independent of the probability of case *j* infecting any other case, the reproductive number for case *j* is a Bernoulli random variable with mean 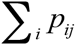. The reproductive number on day *t*, namely *R_t_*, is approximated as the average of the reproductive number of all cases who have symptom onset on day *t*, in which case the mean and standard deviation of *R_t_* are

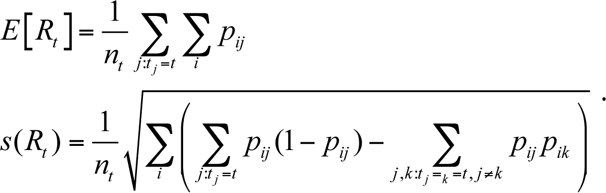

Assuming that *R_t_* is normally distributed, the approximate (1–α)×100% confidence interval is 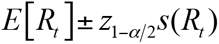.

### Estimation of the serial interval distribution for YF

We assume that the latent period is the same as the incubation period for all human infections of YF. Suppose an infected individual becomes infectious at time 0. Let *t*_1_ be the time at which the infectious individual is bitten by a competent mosquito which becomes infected, *t*_2_ be the time at which this mosquito becomes infectious, and *t*_3_ be the time at which this mosquito bites and infects a human host. The probability distribution function for the serial interval is

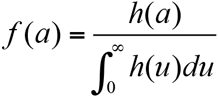

where

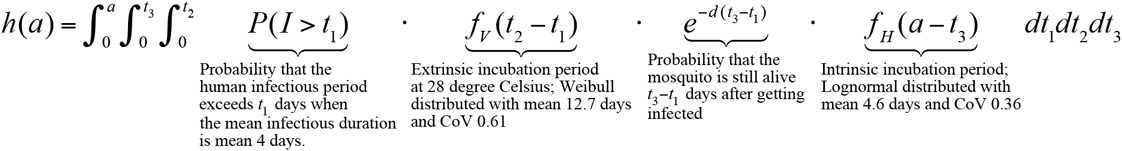

In this calculation, we assume that the infectious period in humans is exponentially distributed with mean 4 days,^29^ and mosquito lifespan is exponentially distributed with mean varying over 1–2 weeks (http://www.dengue.gov.sg/subject.asp?id=12;^17^). We assume that the extrinsic incubation period follows the Weibull distribution with parameters *ν* = 1.7 and *λ_i_* = exp(−7.6 + 0.11*T*) where *T* is the temperature (28 degrees Celsius) as estimated by ref. ^15^ We assume that the intrinsic incubation period follows the lognormal distribution with parameters *μ* = 1.46 and *τ* = 8.1 as estimated by ref. ^15^.

### Dose-response relationship

We assume that vaccine efficacy of *n*-fold fractional-dose vaccines for *n* between 1 and 5 increases linearly with the amount of antigen in the vaccines which is proportional to 1/*n*. In general, if vaccine efficacy of *n*-fold fractional-dose vaccines for *n* between *n*_1_ and *n*_2_ increases linearly with the amount of antigen in the vaccines, then

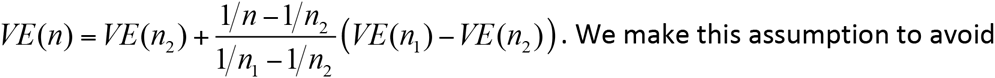

overestimating the benefit of dose fractionation because:

1. If *VE*(5) is at the all-or-nothing benefit threshold, namely *VE*(1)/5, then *VE*(*n*) is also at the benefit threshold (i.e. *VE* (*n*) = *VE* (1)/ *n*) for all *n* between 1 and 5. That is, if five-fold dose fractionation is not beneficial, then dose fractionation is not beneficial for all fractionation below five-fold.
2. The reduction in vaccine efficacy as fractionation increases from 1 is likely to be more gradual than what we have assumed here given that standard dose vaccine efficacy appears to be close to 100%.

Appendix Figure 1 illustrates this dose-response relationship for different values of *VE* (5) with *VE* (1) = 1.

**Appendix Figure 1.**
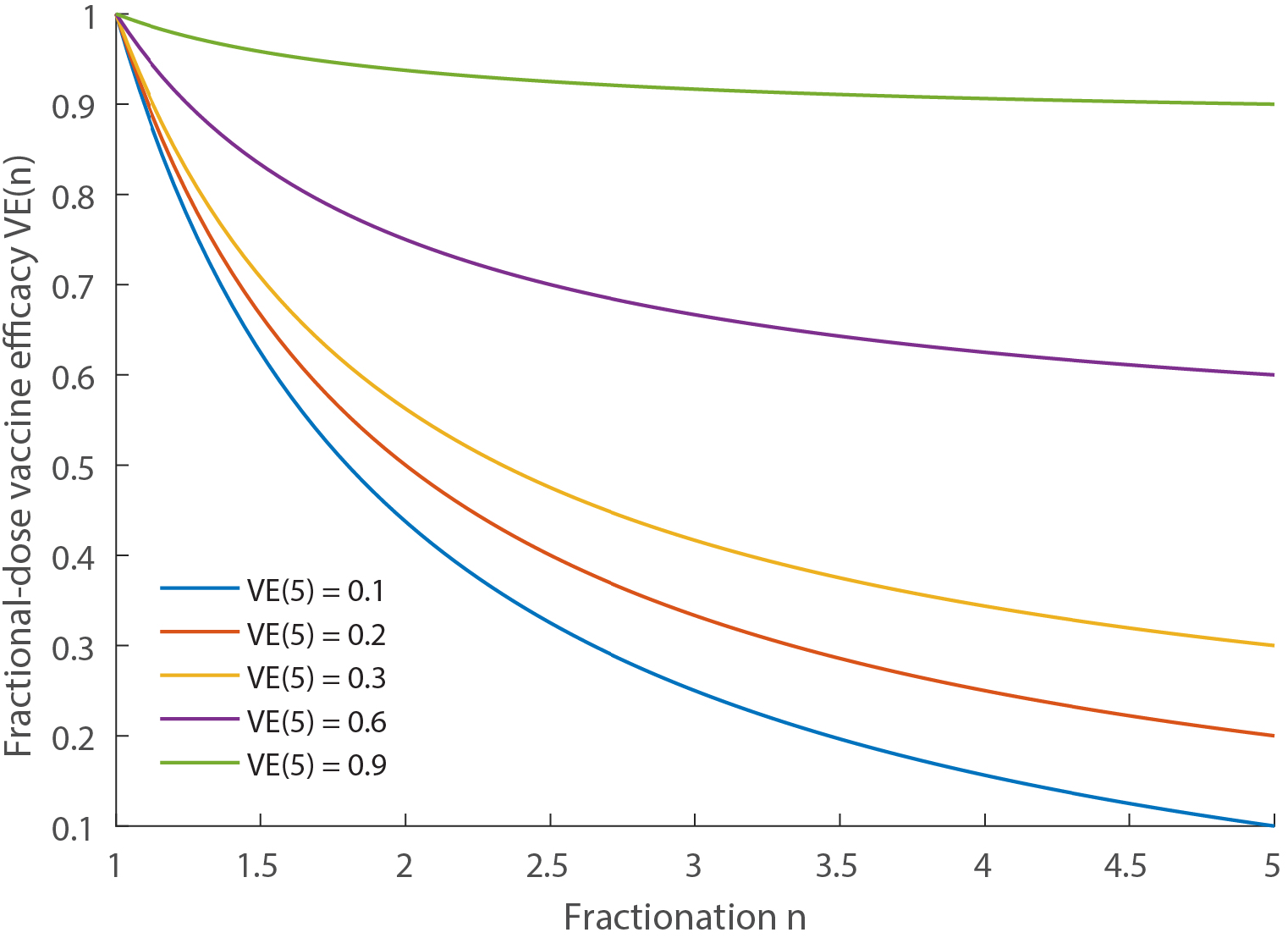
The dose response relationship assumed in the model with *VE*(1) = 1.

### Infection attack rate

We first provide mathematical details on IAR calculations for the case where the population is not stratified into subgroups. If vaccine action is all-or-nothing, then IAR with fractionation n, denoted by *IAR*(*n*), is obtained by solving the equation

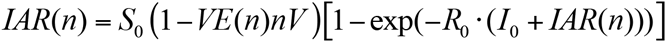

where *R*_0_ is the basic reproductive number, *S*_0_ and *I*_0_ are the initial proportion of population that are susceptible and infectious. As such, dose fractionation reduces IAR if and only if *VE*(*n*) > *VE*(1)/*n*. If vaccine action is leaky, then *IAR*(*n*) is obtained by solving the equation

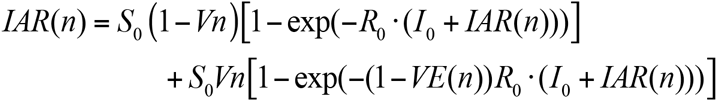

In this case, dose fractionation reduces IAR if and only if

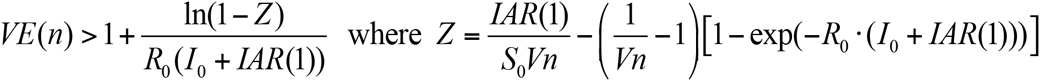

In the special case where *VE* (1) = 1, the benefit threshold can be simplified as

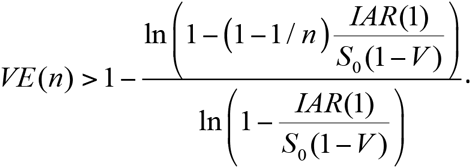

Next, we provide mathematical details on IAR calculations for the general case where there are *m* groups. Let *S_0,i_* and *I_o,i_* be the proportion of susceptible and infectious people in group *i* just before the vaccination campaign begins. Let *V_i_* be the vaccine coverage of standard-dose vaccines for group *i*. If *n_i_* is the fractionation for group *i*, then vaccine coverage of fractional-dose vaccines for group *i* is *V_i_n_i_*. Let 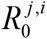 be the expected number of secondary infections in group *j* caused by one infection in group *i* in a completely susceptible population. If vaccine action is all-or-nothing, the group-specific IARs are obtained by solving the equations

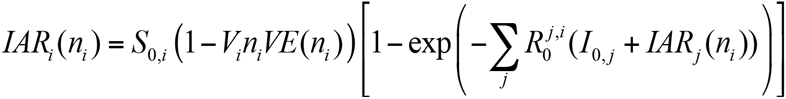

If vaccine action is leaky, then the group-specific IARs are obtained by solving the equations

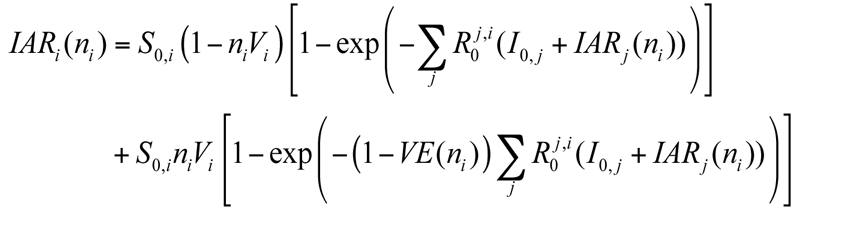

### Heterogeneity in biting rates

A recent study suggested that the mosquito biting rate for individuals aged 20 or above is 1.22 times higher than those age under 20.^24^ To test the robustness of our results against such heterogeneity, we repeat the calculations in Figure 2 and 3 using a model in which the population is stratified with age 20 as the cutoff. For illustration, we use the demographic parameters of Angola where around 55% of the population are under 20. Appendix Figures 2–3 show that our results are unaffected by heterogeneity in biting rates.

**Appendix Figure 2.**
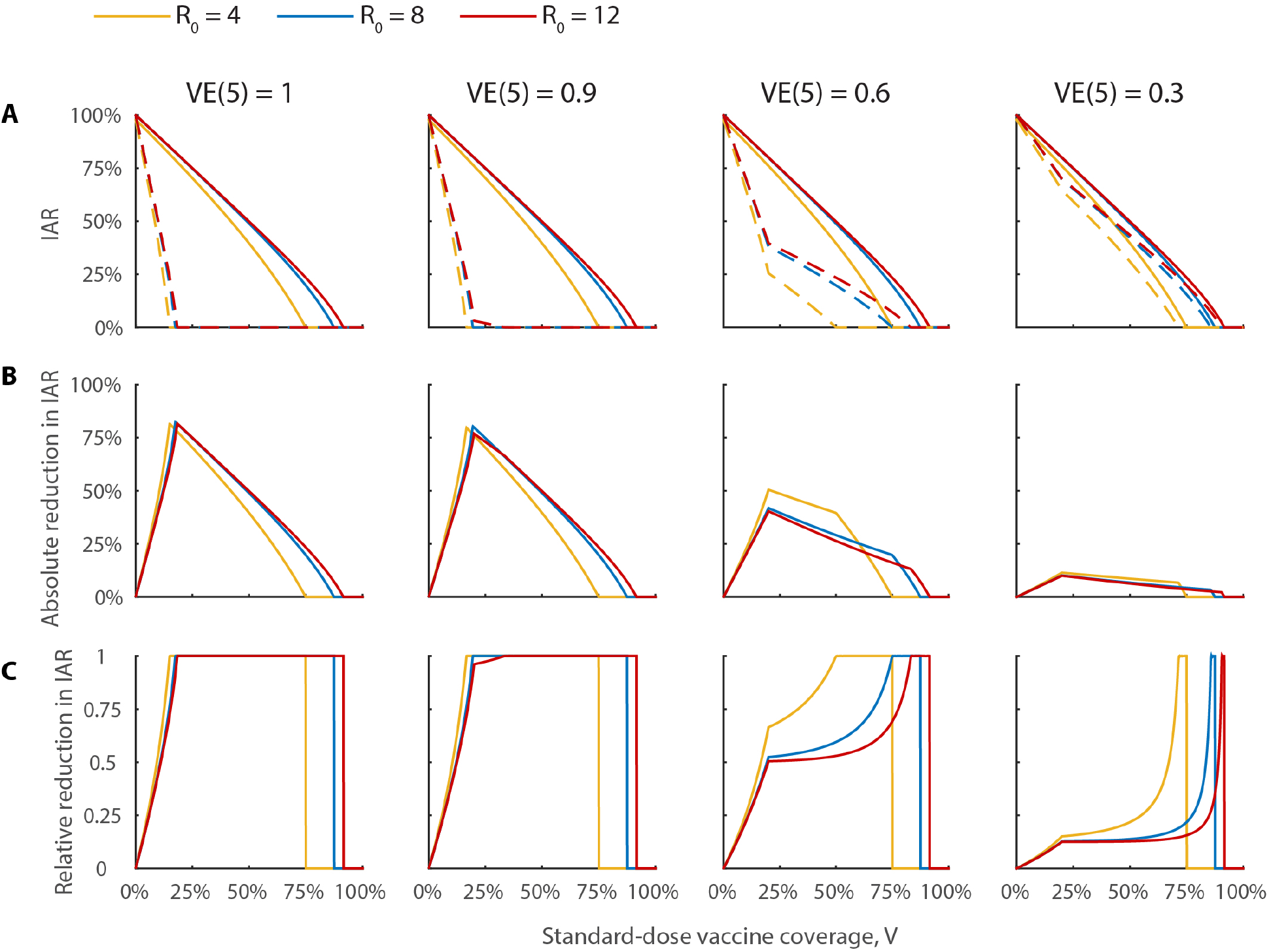
Repeating the calculations in Figure 2 using a 2-age-group model in which those 20 or older were 1.22 times more likely to be bitten by mosquitos compared to those under age 20. A-C. The results are essentially the same as that in Figure 2B–D.

### Vaccine coverage in Angola at the end of 2015

Appendix Table 1 summarizes our estimates of the age distribution and pre-outbreak vaccine coverage in Angola at the end of 2015. The underlying calculations are summarized as follows.

**Appendix Table 1.** Estimated age distribution and vaccine coverage of Angola at the end of 2015.

| Age | Count (100,000) | % of population | Routine immunization coverage (%) | Routine immunization count (100,000) |
| --- | --- | --- | --- | --- |
| 0 - 8 mo | 4.95 | 2.52% | 0% | 0 |
| 9 - 11 mo | 1.65 | 0.84% | 0% | 0 |
| 1 | 6.60 | 3.36% | 72% | 4.752 |
| 2 | 6.60 | 3.36% | 77% | 5.082 |
| 3 | 6.60 | 3.36% | 49% | 3.234 |
| 4 | 6.60 | 3.36% | 63% | 4.158 |
| 5 | 5.50 | 2.80% | 64% | 3.52 |
| 6 | 5.50 | 2.80% | 40% | 2.2 |
| 7 | 5.50 | 2.80% | 40% | 2.2 |
| 8 | 5.50 | 2.80% | 61% | 3.355 |
| 9 | 5.50 | 2.80% | 72% | 3.96 |
| 10 | 4.90 | 2.50% | 43% | 2.107 |
| 11 | 4.90 | 2.50% | 44% | 2.156 |
| 12 | 4.90 | 2.50% | 60% | 2.94 |
| 13 | 4.90 | 2.50% | 52% | 2.548 |
| 14 | 4.90 | 2.50% | 46% | 2.254 |
| 15 | 4.50 | 2.29% | 46% | 2.07 |
| 16 | 4.50 | 2.29% | 50% | 2.25 |
| 17 | 4.50 | 2.29% | 53% | 2.385 |
| 18 | 4.50 | 2.29% | 45% | 2.025 |
| 19 | 4.50 | 2.29% | 37% | 1.665 |
| 20 | 3.70 | 1.89% | 0% | 0 |
| 21 | 3.70 | 1.89% | 0% | 0 |
| 22 | 3.70 | 1.89% | 0% | 0 |
| 23 | 3.70 | 1.89% | 0% | 0 |
| 24 | 3.70 | 1.89% | 0% | 0 |
| 25+ | 70.25 | 35.80% | 0% | 0 |
| Total | 196.25 | 100.00% | - | 54.861 |

*Age distribution*. The age distribution of Angola with 5-year age bands is obtained from The World Factbook at the CIA (https://www.cia.gov/library/publications/the-world-factbook/graphics/population/AO_popgraph%202015.bmp). We assume that the age distribution within each 5-year age band is uniform.

*Pre-outbreak vaccine coverage*. Routine YF vaccination in Angola began in 1997. We obtain the annual routine immunization coverage among children aged 12–23 months between 1997 and 2015 from the WHO/UNICEF immunization estimates (http://apps.who.int/immunization_monitoring/globalsummary/estimates?c=AGO).

### Kinshasa vaccination campaign in July-August 2016

Appendix Table 2 summarizes our estimates of the age distribution and pre-campaign vaccine coverage in Kinshasa at the end of June 2016. The underlying calculations are summarized as follows.

*Age distribution*. The age distribution of Kinshasa with 5-year age bands is obtained from CIA Facts (https://www.cia.gov/library/publications/the-world-factbook/graphics/population/CG_popgraph%202015.bmp). We assume that the age distribution within each 5-year age band is uniform.

*Pre-campaign vaccine coverage*. YF vaccine coverage in Kinshasa at the end of June 2016 comprises the following two components:

1. Routine YF childhood immunization which started in 2004. We obtain the annual routine immunization coverage among children aged 12–23 months between 2003 and 2015 from the WHO/UNICEF immunization estimates (http://apps.who.int/immunization_monitoring/globalsummary/estimates?c=COD). We assume that routine immunization coverage between age 9 months and 1 year at the end of June 2016 is 0 because of the YF severe vaccine supply shortage in 2016.
2. Emergency vaccination of around 1 million people in Kinshasa during May-June 2016 (ref ^25^; personal communication, Bruce Aylward and Alejandro Costa, WHO). We assume that the coverage of this emergency vaccination was uniformly distributed across age.

The WHO has planned to deliver around 2.5 million standard-dose vaccines in the Kinshasa vaccination campaign in July-August 2016 (personal communication, Bruce Aylward and Alejandro Costa, WHO). The target population size in Kinshasa is around 8.05 million, with potentially 1–2 million additional people from adjacent areas. Of the 2.5 million standard-dose vaccines, 200,000 will be administered to children age 9 months to 2 years in standard dose (which is sufficient for vaccinating all unvaccinated children in this age range) and the rest of the vaccine stockpile will be administered to the rest of the population as five-fold fractional-dose vaccines. As such, when considering alternative age cutoff for fractional-dose vaccines, we assume that the vaccine stockpile is sufficient to vaccinate all individuals in Kinshasa that have not yet been vaccinated as of 30 June 2016.

### Leaky versus All-or-Nothing YF Vaccine Action

We consider the all-or-nothing YF vaccine action mechanism more likely than the leaky vaccine action model for the following reasons: based on the limited evidence on immunogenicity of fractional doses to date, we consider it unlikely that reducing the dose five-fold or perhaps further from current preparations would result in dramatically lower efficacy of the leaky type. Visual inspection of the data from a dose fractionation trial of the 17DD vaccine in Brazil shows that for doses down to 47x below the standard dose, the distribution of serologic responses was indistinguishable from those for the standard dose, suggesting that efficacy should be nearly equivalent to that for full doses. This was confirmed by the analysis of peak viremia, which was equivalent for standard dose and for doses down to 11% of the full dose (9-fold fractionation). It was further confirmed by peak cytokine responses, which were comparable to the standard dose for all cytokines tested, down to at least a 9-fold fractional dose. For even lower doses, the proportion seroconverting after vaccination was lower than the 97% observed for the full dose, but the antibody response among the seroconverters appears to be similar at all doses.^9^ These data collectively suggest that down to approximately 9-fold fractional dosing of this vaccine the response should be equivalent, and that for further fractionation there may be a failure to induce any substantial response in a fraction of recipients, but the neutralizing antibody titres in those who do respond should be comparable. This pattern is consistent with an all-or-nothing model.

**Appendix Figure 3.**
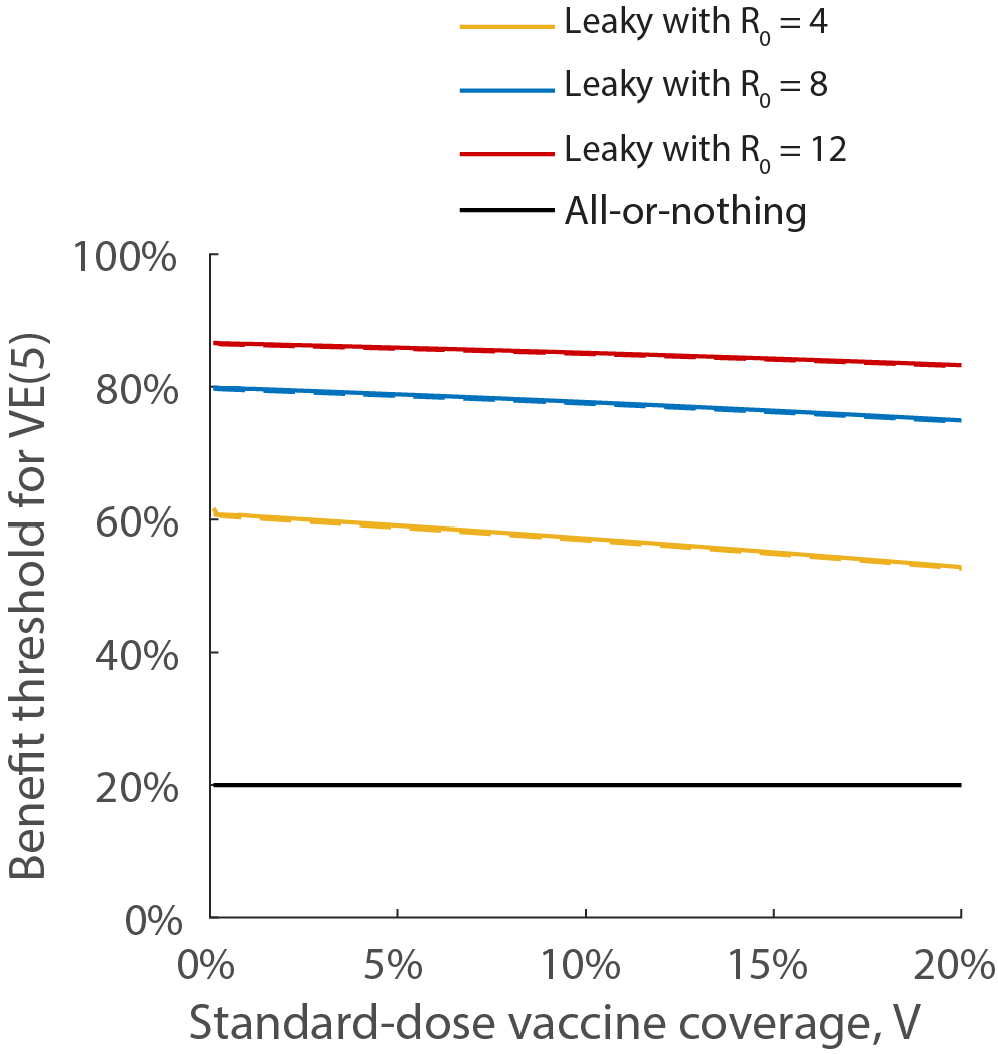
Repeating the calculations in Figure 3 using a 2-age-group model in which those 20 or older were 1.22 times more likely to be bitten by mosquitos compared to those under age 20. The solid and dashed curves show the results without and with age stratification, respectively.

